# Newborns modulate and synchronise their Non-Nutritive Sucking behaviour to the temporal structure of sound

**DOI:** 10.64898/2026.10.06.756750

**Authors:** Martina Arioli, Alessandra Federici, Valentina Silvestri, Matteo Santini, Lorenzo Colombo, Maria Lorella Giannì, Davide Bottari, Viola Macchi Cassia

**Affiliations:** Department of Psychology, University of Milano-Bicocca, Milan, Italy; MoMiLab, IMT School of Advanced Studies Lucca, Lucca, Italy; Faculty of Experimental Psychology Helmholtz Institute, Utrecht University. Utrecht, Netherlands; Department of Clinical Sciences and Community Health, University of Milano, Milan, Italy; Fondazione IRCCS Ca’ Granda Ospedale Maggiore Policlinico, NICU, Milan, Italy

**Keywords:** Newborns, Non-Nutritive Sucking, Sensori-Motor Synchronization, Music, Coherence

## Abstract

Sensorimotor synchronization, the ability to align motor behaviour with the temporal structure of external stimuli, is thought to emerge gradually during development, but its origins remain unclear. Here, we investigated whether newborns can adapt their sucking behaviour and synchronise it to the temporal structure of auditory stimuli within the first days of life. Forty-one healthy full-term newborns (24-72 hours old) were exposed to two three-minute auditory streams differing in temporal structure: a highly structured stream with a stable temporal pattern and a highly variant stream with greater temporal variability, while their non-nutritive sucking (NNS) was continuously recorded. We assessed changes in NNS frequency and pressure and quantified NNS-sound coupling using magnitude-squared coherence at frequencies distinctive to each auditory stream. Structured stimulation promoted a more organised NNS, with more suck bursts than the variant stream, and increased sucking vigour, with a transiently higher suck peak rate during the first minute relative to the variant stream and greater suck pressure relative to silence. Coherence analyses revealed that coherence was selectively higher at the frequencies distinctive to each auditory stream, both for the structured and the highly variant stimulus. These findings show that, within hours of birth, NNS behaviour can selectively align with stimulus-specific temporal regularities, even when these are weakly predictable and lack a dominant periodicity. This early capacity to extract temporal structure and coordinate motor behaviour may provide a foundation for the subsequent development of sensorimotor synchronisation.

**Highlights:**

- Structured sounds elicit more organized and frequent sucking compared to variant sounds.
- NNS selectively couples to stimulus-specific temporal frequencies.
- NNS synchronization also occurs for temporally variable auditory stimuli.
- Sensorimotor synchronization is evident within hours of birth.

## 1. Introduction

The ability to adapt or synchronize one’s motor rhythm to the rhythm of an external stimulus represents a fundamental mechanism for early communicative development, supporting temporal coordination during caregiver-infant interactions and facilitating processes such as turn-taking and the extraction of temporal regularities relevant for language learning (Provasi et al., 2014; Tillman et al., 2026; Woodruff Carr et al., 2014). Although existing literature has documented the presence of this ability very early in development, it remains unclear whether it emerges through postnatal auditory-motor experience or whether it is already shaped by prenatal factors, either through the rich acoustic and vestibular rhythmic stimulation experienced in utero or through intrinsic self-generated rhythmic activity. Testing this capacity at the very onset of postnatal life, before any substantial postnatal multisensory experience has occurred, offers a unique opportunity to disentangle these possibilities. The present study aims to address this gap by investigating early motor adaptation and synchronization to external auditory rhythm in 24-72-hour-old newborns.

Already during intrauterine life, the foetus generates a rich repertoire of rhythmic activity, spanning both physiological processes (e.g., cardiac pulsations and respiratory movements) and motor behaviours. From approximately the 13th to the 16th week of gestation, rhythmic patterns of flexion and extension can be observed in the foetus, sometimes coordinated with arm movements (Reinold, 1979). Among these early motor rhythms, Non-Nutritive Sucking (NNS) is particularly well characterized: it emerges around the 10th week of gestation and becomes progressively more organized as gestation advances. By term, it reaches a structured pattern of suck bursts separated by pauses, occurring at a frequency of approximately 1 Hz, comparable to that observed in newborns (Popescu et al, 2008; Wolff, 1968).

Rhythmic activities that are part of the human behavioural repertoire across the lifespan are characterized by a specific tempo, referred to as the spontaneous motor tempo, which can be modified in response to environmental factors. In particular, the ability to adapt or synchronise an individual’s motor rhythm to the rhythm of an external stimulus is known as sensorimotor synchronization (SMS), a process thought to support the temporal coordination of behaviour during social interactions and to contribute to language-related development (e.g., Provasi et al., 2014). Indeed, it has been proposed that impairments in SMS, together with difficulties in perceptual timing, may serve as markers for identifying children at risk for speech and language disorders (Atypical Rhythm Risk Hypothesis; Ladányi et al., 2020).

SMS develops gradually, with significant improvements continuing well into adulthood (Drake et al., 2000; Thompson et al., 2015). Within SMS, a distinction can be drawn between tempo flexibility and precise synchronization. The latter-the ability to precisely align a motor response with the temporal structure of an external rhythmic stimulus-can be reliably observed only from around 2.5 years (e.g., Rocha et al., 2022) and continues to refine across development. Before this age, SMS is instead visible only in terms of tempo flexibility, namely the ability to adjust movement speed in response to faster or slower auditory tempo (Zentner & Eerola, 2010). For instance, Rocha et al. (2021) showed that infants aged 5–11 months could not reliably synchronise their movements to auditory rhythms but, by 11 months, could slow their spontaneous motor tempo toward the auditory tempo. Similarly, Nguyen et al. (2025) found evidence of coarse auditory-motor coupling in 3-, 6-, and 12-month-old infants, as their spontaneous movements were better predicted by the temporal envelope of the original than temporally shuffled music.

In neonates, tempo flexibility has been investigated mainly using NNS recordings. Using tactile stimulation, Poore and colleagues (2008) showed that administering preterm infants with patterned orocutaneous input, in the form of regular pneumatic pulse trains delivered through a silicone pacifier, improved the regularity and coordination of the NNS behaviour. Using auditory stimulation, Bobin-Bègue and colleagues (2006) reported that exposing newborns to a sound accelerated by 15% relative to their previously recorded NNS spontaneous motor tempo resulted in an acceleration of the NNS rate. However, no corresponding deceleration was observed when the sound was slowed down. Using a different paradigm, Zimmerman and Foran (2017) exposed older infants (1 to 6 months) to fixed auditory frequencies (1, 2, and 4 Hz) rather than tempi individually calibrated to each infant’s spontaneous motor tempo, and found a deceleration of NNS in response to the 1- and 2-Hz stimuli, but not to the 4-Hz stimulus. This decrease in NNS rate may reflect attentional capture (e.g., Valenza et al., 1994) elicited by frequencies closer to the infants’ sucking spontaneous motor tempo (1 and 2 Hz), an effect that did not emerge with the faster, less proximal stimulus frequency.

Together with the developmental trajectory outlined above, these findings suggest that external rhythmic stimuli promote motor adaptation from the very beginning of postnatal life. However, it remains unknown whether newborns right after birth can actually synchronise their motor behaviour with the temporal structure of an external auditory stimulus. Previous studies have primarily measured changes in overall sucking rate in response to rhythmic stimulation (e.g., Valenza et al., 1994), which cannot distinguish genuine stimulus-specific coupling from nonspecific effects of attention, arousal, or behavioural activation. In particular, NNS studies have not complemented changes in sucking frequency with established measures of sensorimotor synchronization, such as phase-locking indices, that are routinely employed in the motor entrainment literature. A further limitation is that previous studies have relied mainly on simple isochronous pulse trains, leaving open whether newborns can track the temporal organization of more complex and variable auditory sequences. Given that non-nutritive sucking (NNS) is already highly organized at birth it may provide a uniquely sensitive behavioural window onto the emergence of sensorimotor synchronization.

The present study aims to address these gaps by investigating neonatal NNS motor adaptation (i.e., changes in sucking frequency and/or pressure) and sensorimotor synchronization (i.e., the coupling between the NNS signal and the temporal structure of the auditory stimuli) in response to musical stimuli varying in their temporal structure. We recorded the NNS behaviour of 24-to 72-hour-old newborns while they were exposed to two auditory streams: one characterized by a high degree of temporal structure and one by a highly variable temporal structure. As a first step, and in line with previous rate-based approaches, we examined differences in NNS frequency and pressure across conditions, hypothesising that newborns would show increased NNS in response to the structured compared to the variant stream. Second, to test whether sucking behaviour became synchronised to the two auditory streams, we measured magnitude-squared coherence between the NNS signal and each auditory stimulus at their distinctive frequencies. Unlike previous rate-based approaches, which compared changes in NNS rate to a single fixed frequency defined by an isochronous stimulus, this method quantifies coupling across the full range of frequencies present in the auditory stream, allowing us to test whether the newborns’ sucking selectively tracks the specific temporal structure of each auditory stream, rather than responding to auditory stimulation in a generic, undifferentiated way.

## 2. Method

### 2.1. Participants

The final sample comprised 41 healthy, full-term newborns (21 females; mean gestational age: 38.7 weeks, range 37-41 weeks; mean age = 50.09 hours, range = 23.27-88.97 hours; mean weight: 3255 g, range 2300-4400 g). An additional 19 participants were tested but later excluded due to drowsiness/fussiness (n = 13), technical errors (n = 3), or refusal of the pacifier (n = 3). Sample size was not determined by an a priori power analysis; rather, we aimed to recruit a larger sample than previous studies investigating newborns’ attention to auditory stimuli using a similar design (e.g., N=12; Standley, 2000), in order to mitigate potential power issues. To further validate this, we performed a sensitivity analysis using the SIMR package for R (Green & Macleod, 2016) to determine the smallest detectable effect with 80% power and alpha = .05 using mixed effect models, with 1000 simulations. This was not an observed/retrospective power analysis; rather, we systematically varied the effect sizes of the Condition fixed effects of interest for our hypothesis (Perugini et al., 2018). Results confirmed that our sample of 41 participants (contributing 105 observations) was sufficient to detect a difference of 1.07 mbar of pressure, 1.2 bursts/minute and 9.2 peaks/minute. Newborns were recruited from the maternity ward of the Mangiagalli Clinic in Milan. All participants met the following inclusion criteria: gestational age greater than 37 weeks, birth weight above 2300 g, an Apgar score of at least 9 at 5 minutes and normal hearing. Testing was conducted while the infants were awake and alert. All participants were from middle-class families, with parents self-identifying as White. Written informed consent was obtained from the parents prior to testing. The study protocol adhered to the ethical standards of the Declaration of Helsinki (BMJ 1991; 302:1194) and was approved by the Comitato Etico Milano Area 2; ID: 694; Approval N. 952_2021.

### 2.2. Stimuli

The stimuli consisted of two auditory streams created by combining piano, mallet, and vibraphone sounds using Logic Pro 10.5. The two streams were designed to differ in temporal structure. The structured stream consisted of a 3-minute repetition of a 2-second basic sequence, which was a modified version of the one used by Winkler and colleagues (2009) in their study on newborns’ detection of beat omissions within complex auditory sequences. It maintained the same metric structure while using different sounds. The variant stream was created by generating four 2-second base sequences derived from the 2-second base sequence of the structured stream, with both the order of the notes and the intervals between them randomly shuffled. Importantly, each of the 2-second sequences contained the same type, and number of notes as the 2-second base sequence of the structured stimulus, ensuring full comparability in low-level acoustic features. The sequences were then randomly concatenated to create a continuous 1-minute stream, with the constraint that the same sequence could not be repeated consecutively more than twice. This 1-minute stream was then repeated three times to obtain the final 3-minute stimulus (Figure 1a). The stimuli can be accessed at the following link: https://osf.io/3dkxc/overview?view_only=8b740e982f8843d1843499e8af0dbac4

**Figure 1.**
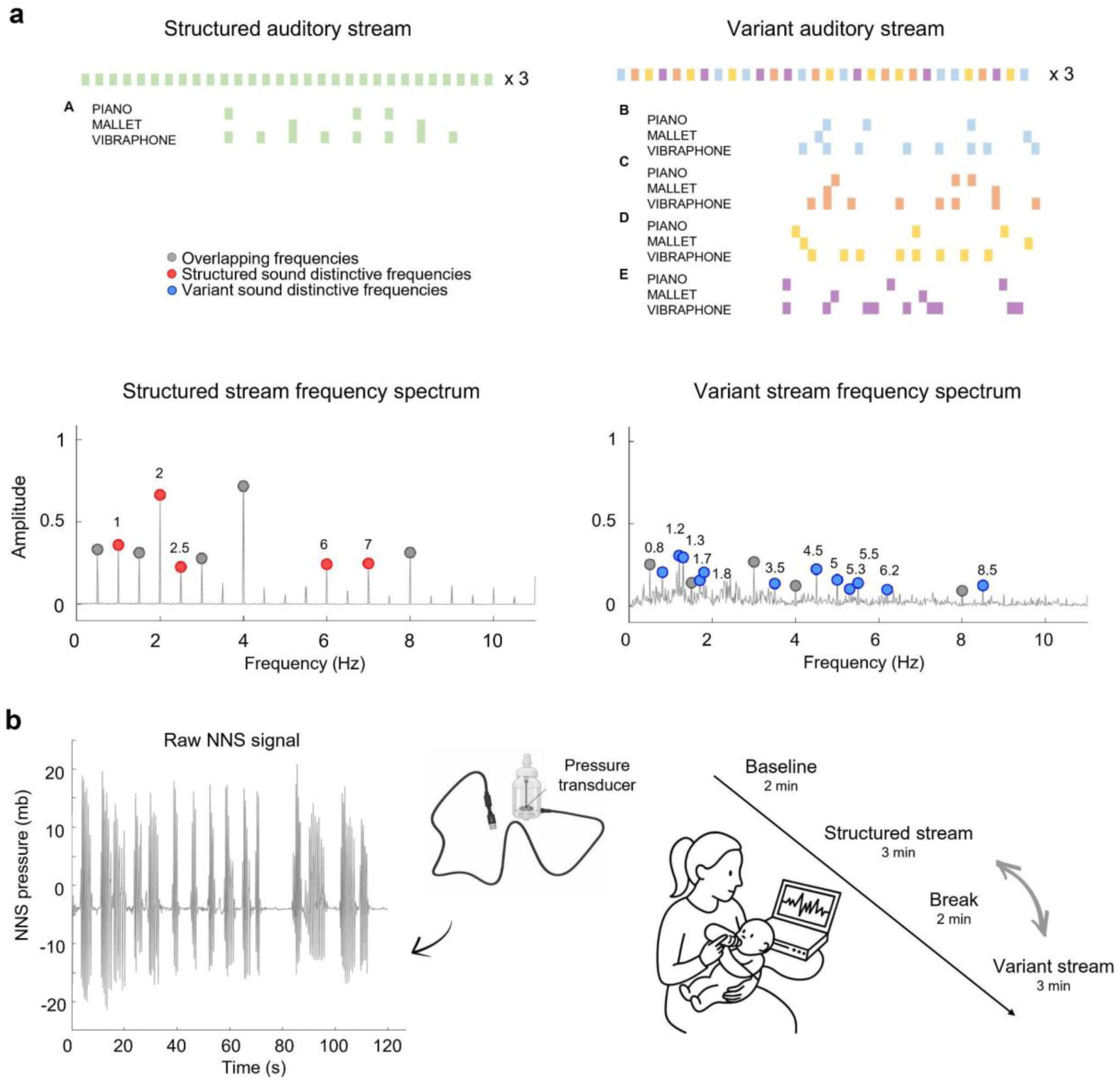
Panel a provides an overview of the highly structured and highly variant auditory streams and their spectral characteristics. The upper boxes illustrate the combination of 2-s sequences presented during the first minute for each stream (structured on the left and variant on the right), which was repeated three times to obtain the 3-min stimulation for each condition. The lower diagrams show the power spectra of the first minute of the structured and variant auditory streams, respectively. Overlapping frequencies between the two streams are marked in grey; distinctive frequencies in each stream are marked in red (Structured stream) and blue (Variant stream). Panel b illustrates an exemplar NNS (Non-Nutritive Sucking) signal and the apparatus and procedure, including the custom-made pacifier with a pressure transducer used for NNS data collection.

### 2.3. Apparatus and procedure

Infants were tested when in alert state, seated on the experimenter’s lap in a semi-inclined position throughout the experiment. NNS behaviour was recorded at 166.7 Hz using a feeding bottle fitted with a custom-made pacifier containing an embedded pressure transducer connected to a computer (Figure 1b). The experimenter encouraged the infant to take the pacifier by gently stroking the infant’s cheek, while holding it attached to the recording apparatus. Data acquisition began as soon as the infant had the pacifier in his/her mouth. Each pacifier was used only once per participant and was discarded after each session. The researcher holding the newborn was instructed not to engage or look at the infant during the testing, and to keep their head tilted backward throughout the session, so as not to be seen by the infant.

A second researcher was present in the room and operated the data acquisition system, ensuring that the apparatus functioned properly and that data were correctly recorded. One or both parents remained in the room for the entire experimental session, seated behind the experimenter and outside the infant’s direct line of sight. Parents were instructed to interact with the infant only if signs of discomfort were observed, in which case the experimental session was interrupted. Room lighting was kept dim throughout the experiment.

Auditory streams were presented in free field at an intensity of 70 dB through the computer’s loudspeakers, positioned approximately 40 cm from the infant. Each experimental session began with a 2-minute silent baseline period, followed by 3 minutes of acoustic stimulation with either the structured or the variant stream, with condition order counterbalanced across participants. This was followed by an additional 2-minute break period to separate the two stimulation blocks, and then by a further 3 minutes of acoustic stimulation with the other stream (Figure 1b). The total duration of each session was 10 minutes.

### 2.4. Data reduction

#### 2.4.1. NNS frequency and pressure

The raw suck pressure signal was processed in MATLAB (MathWorks, Natick, MA; version 2022b). Preprocessing was performed based on the procedure described by Barlow et al. (2014, 2023). First, the NNS signal was downsampled to 100 Hz. It was then band-pass filtered (0.5-20 Hz) to remove low-frequency offsets related to tongue and jaw posturing, thermal drift resulting from contact between the oral apparatus and the pacifier bulb, and high-frequency noise. Finally, the signal was smoothed using LOWESS over 16 data points (approximately 160 ms).

NNS suck peaks and bursts were identified using the *findpeaks* function in MATLAB. A suck peak was defined as a sucking event in which the pressure exceeded 1.5 mb (Barlow et al., 2014, 2023; Liao et al., 2019); a suck burst was defined as two or more suck peaks separated by an inter-peak interval of less than 1.2 s (Barlow et al., 2023). The dependent variables extracted from the signal included, for each sound condition (baseline, structured stream, variant stream), the mean number of suck peaks per minute, mean number of bursts per minute, mean burst duration (s), mean number of suck peaks per burst, interburst distance (s), and mean suck pressure (mb). To assess whether auditory stimulation elicited an initial freezing of sucking rate, the mean number of suck peaks was also extracted for the first minute of stimulation, separately for the structured and variant conditions.

#### 2.4.2. NNS-sound coherence at sound-distinctive frequencies

The two sound streams, originally sampled at 48.000 Hz, were imported and downsampled by a factor of 10. To identify the distinctive frequencies carrying the temporal structure of each stream, we computed the power spectrum in the 0-10 Hz range using a Fourier transform (FFT) on the envelope of a 1-minute-long segment, corresponding to the minimum duration required for the variant stream to contain its full variability. To retain only clearly distinguishable spectral components and reduce the detection of spurious peaks, prominent spectral peaks were identified using *findpeaks* with the minimum peak prominence set to 30% of the maximum spectral amplitude recorded in the signal (Figure 1b). Mutuating the analytical framework from frequency tagging approaches in electrophysiological research, overlapping frequencies between the selected spectral peaks of the two streams were then discarded in order to isolate stream-specific frequencies (see Rossion et al., 2015 and Norcia et al., 2015). From this sample of selected frequencies only the frequencies with 0.1 Hz resolution were finally kept for the coherence analysis in order to match the frequency resolution of the NNS-sound coherence analysis. As a result, five distinctive frequencies were identified for the structured stream (i.e. 1, 2, 2.5, 6, 7 Hz) and twelve distinctive frequencies were identified for the variant streams (i.e., 0.8, 1.2, 1.3, 1.7, 1.8, 3.5, 4.5, 5, 5.3, 5.5, 6.2, 8.5 Hz) (Figure 1b). Notably, the distinctive frequencies of the structured and variant sound streams fell within a similar range, reducing the likelihood that differences in NNS-sound coherence were driven by the endogenous NNS spectrum.

To compute NNS-sound coherence at sound distinctive frequencies, the auditory streams (highly structured and highly variant) were first imported at their native sampling rate and converted to mono by averaging the two stereo channels. The NNS signal was baseline-corrected by subtracting the individual median pressure value computed across the entire recording session, to account for inter-individual differences in sucking pressure. Then, the NNS signal of each sound condition was temporally aligned to the corresponding auditory stimulus onset. Because of a small delay (≈3 s) between the sound onset and the start of the signal recording, NNS and sound signals were temporally realigned using the recorded onset timestamps. Only the overlapping signal was retained for subsequent analyses, and all recordings were subsequently trimmed to 176 s, corresponding to the minimum available recording duration across participants. For coherence analysis, the NNS signal was interpolated to match the sampling rate of the auditory stimulus. Both NNS and sound signals were then normalized using z-score transformation and downsampled by a factor of 10, resulting in a common sampling rate of 4.800 Hz. The amplitude envelopes of the sucking and sound signals were then extracted using the Hilbert transform. The Direct Current (DC) offsets were removed from both envelopes, which were subsequently low-pass filtered at 10 Hz using a third-order zero-phase Butterworth filter to isolate slow amplitude modulations relevant for rhythmic coupling. Magnitude-squared coherence between the NNS and sound envelopes was computed using the *mscoherence* function in MATLAB. Hereafter, this measure is referred to as coherence. Coherence values were estimated for each participant and auditory stream condition (structured, variant) using a 18-s Hamming window with 50% (9 s) overlap, within the 0.1-10 Hz frequency range, and with a frequency resolution of 0.1 Hz. To explore whether coherence values changed over time, coherence was also computed separately for each minute (1st, 2nd, 3rd) in each condition using the same parameters (i.e. 58.67 s, corresponding to one third of the 176-s trimmed signal).

## 3. Results

### 3.1. NNS frequency and pressure

To test whether NNS frequency and pressure varied between conditions, we fitted a linear mixed model for each of the dependent variables extracted with sound condition (baseline, structured, variant) as fixed factor and a random intercept for participants. The marginal ΔR² was used as the effect size for the fixed effects in the linear mixed-effects models. It was calculated as the difference in marginal R² between the full model and the corresponding reduced model without the effect of interest. Results showed that the mean number of suck peaks per minute did not vary across sound conditions, *F*(2,66.04) = 2.47, *p* = .09, but when the same model was fitted for the number of peaks in the first minute of acoustic stimulation (instead of the average across the 3 minutes of stimulation) the condition effect turned to be significant, *F*(2,67.17) = 4.91, *p* = .01, ΔR² =.047, with the number of peaks in the first minute being significantly higher during the structured sound listening (*M* = 31.4, *SE* = 4) compared to the variant sound listening (*M* = 18.8, *SE* = 3.67), *t*(66.1) = 3.1, *p* = .009. Neither condition differed significantly from baseline (all *p*s > .1) (Figure 2a).

**Figure 2.**
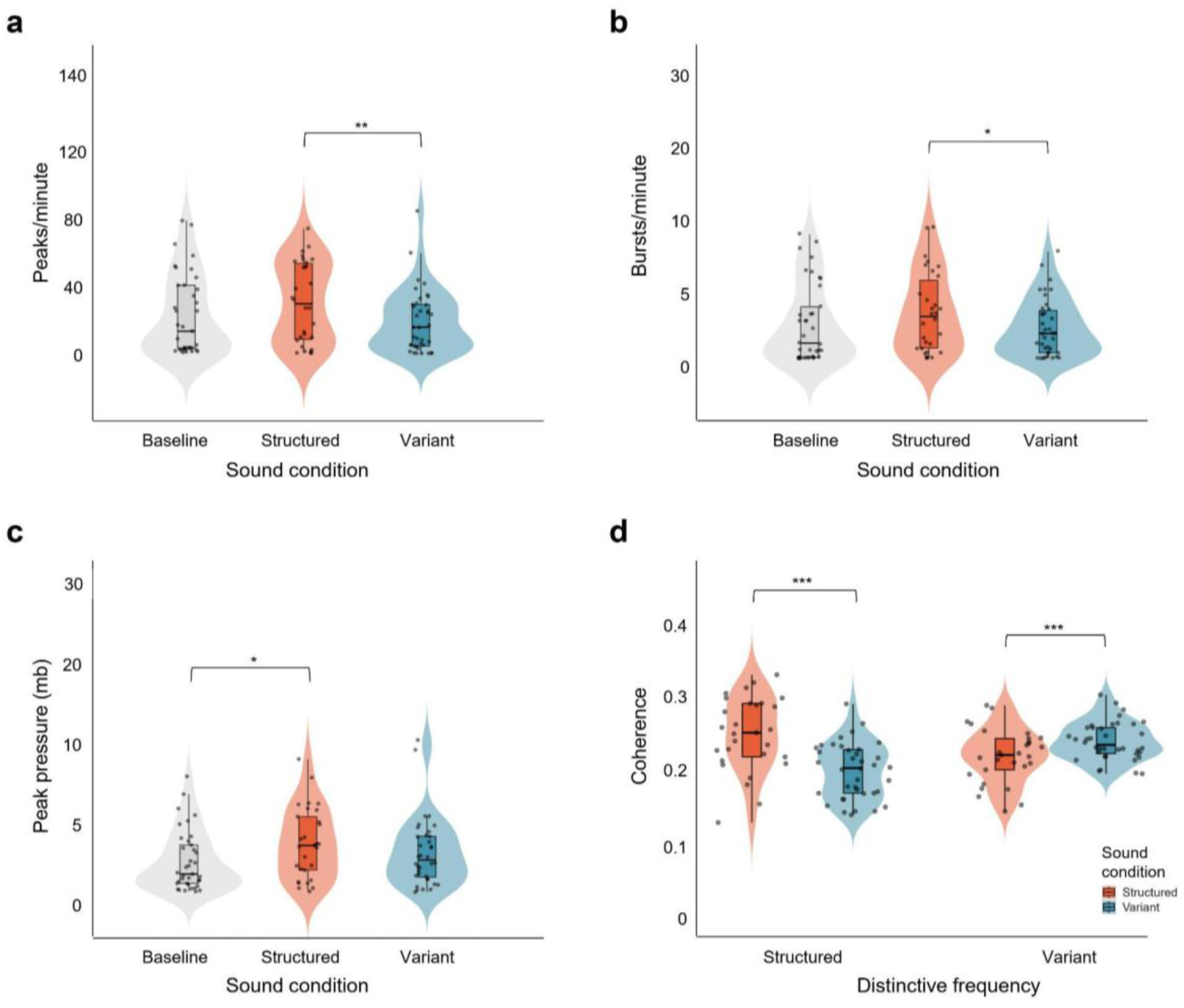
Number of suck peaks in the first minute of structured and variant auditory stimulation compared with mean baseline peaks (panel a), number of bursts per minute across conditions (panel b), mean suck pressure across conditions (panel c), and coherence values between NNS and the structured and variant acoustic streams at the distinctive frequencies of each sound (panel d). Violin plots depict the data distribution (kernel density), with jittered points representing individual observations. Embedded boxplots show the median (central line) and interquartile range (25th-75th percentiles; box edges), with whiskers extending to 1.5 x the IQR. * *p* < .05; ** *p* < .01; *** *p* < .001.

The number of suck bursts per minute varied across sound conditions, F(2,65.36) = 4.46, *p* = .015, ΔR² = .035. Specifically, the number of bursts in the structured condition (*M* = 3.13, *SE* = .43) was higher compared to the variant condition (*M* = 1.99, *SE* = .4), *t*(65.1) = 2.9, *p* = .015; neither the structured nor the variant condition differed significantly from baseline (all *p*s > .09, Figure 2b). No differences were found for the burst duration (s), number of peaks per burst and interburst distance (s) (all *p*s > .2).

In line with the burst results, results also showed that the mean suck pressure varied across conditions, *F*(2,63.43) = 5.43, *p* = .006, ΔR² = .04, with the structured auditory streams (*M* = 3.99 mbar, *SE* = .39) eliciting higher suck pressure than the baseline (*M* = 2.87 mbar, *SE* = .36), *t*(62.9) = 3.21, *p* = .006 (Figure 2c).

### 3.2. NNS-sound coherence at sound-distinctive frequencies

To test whether coherence values to the distinctive frequencies of each sound stream differed between the two sound conditions (structured vs variant), we ran a linear mixed model with coherence values as dependent variable, distinctive frequencies (structured vs variant), sound condition (structured vs variant) and their interactions as independent variables, and random intercept for participants.

Results showed a significant main effect of sound condition, *F*(1,1012.4) = 5.99, *p* = .01, ΔR² = .01, indicating that overall coherence was higher during the structured sound listening (*M* = .08, *SE* = .003) compared to the variant sound (*M* = .07, *SE* = .003). Importantly, the Distinctive Frequencies × Sound condition interaction was also significant, *F*(1,1093.3) = 12.59, *p* < .001, ΔR² = .01. Bonferroni corrected simple effects revealed that during the structured sound listening coherence values were higher for the structured distinctive frequencies (*M* = .09, *SE* = .005) compared to the variant distinctive frequencies (*M* = .07, *SE* = .003), *t*(1099) = 2.79, *p* = .005; conversely, during the listening of the variant acoustic stream, coherence values for the variant distinctive frequencies (*M* = .07, *SE* = .003) were higher than coherence to the structured distinctive frequencies (*M* = .06, *SE* = .004) , *t*(1099) = 2.2, *p* = .03.

This pattern of results was confirmed by a further analysis including minute (1st, 2nd, 3rd) as a fixed effect, both as a main effect and in interaction with the other factors. While there was no significant effect of minute (all *p*s > .08), the analysis showed a Sound condition main effect, *F*(1,2282.6) = 5.78, *p* = .02, ΔR² = .011, with higher NNS-sound coherence during the listening of the structured (*M* = .25, *SE* = .005) compared to the variant (*M* = .23, *SE =* .005) acoustic stream, as well as a Distinctive Frequencies x Sound condition interaction *F*(1,3369.7) = 32.57, *p* < .001, ΔR² = .01. Bonferroni corrected simple effects showed that, during the structured sound listening, coherence was higher for the distinctive structured frequencies (*M* = .26, *SE* = .008) than for the distinctive variant frequencies (*M* = .23, *SE* = .006), *t*(3368) = 3.221, *p* = .001. Conversely, during the variant sound listening, coherence was higher for the distinctive variant frequencies (*M* = .25, *SE* = .005) than for the distinctive structured frequencies (*M* = .21, *SE* = .007), *t*(3368) = 4.99, *p* < .001 (Figure 2d).

## Discussion

The present study investigated newborns’ ability to adapt and synchronise their non-nutritive sucking (NNS) behaviour to the rhythm of external auditory stimuli, specifically a highly structured and a highly variant auditory stream. Overall, results showed that within hours from birth, NNS behaviour is not only modulated in its overall rate and pressure by auditory input, but also becomes selectively coupled to the distinctive frequencies of the presented auditory streams.

Newborns produced more suck bursts while listening to the structured compared to the variant stream. Since a burst is a cluster of temporally close suck peaks, an increase in burst rate indicates that a larger proportion of sucking activity was organized into such clusters, rather than occurring as isolated, unclustered cycles. This suggests that exposure to a temporally regular auditory input supported the organization of sucking behaviour into more structured rhythmic patterns. This is consistent with previous evidence that patterned, regular stimulation can enhance the organization and coordination of NNS in preterm infants, where regular pneumatic pulse trains delivered through a silicone pacifier improved suck burst regularity (Poore et al., 2008). The present findings extend this evidence along three dimensions: from preterm to full-term neonates, from tactile to auditory stimulation, and from simple, isochronous pulse trains to complex, naturalistic auditory sequences, suggesting that the facilitatory effect of temporal regularity on NNS organization in newborns generalizes across sensory modalities and stimulus complexity.

While the burst results point to a sustained organizational effect of acoustic temporal regularity, the analysis of suck peak rate reveals a complementary, more time-limited signature of this effect. The number of suck peaks per minute did not differ between conditions when averaged across the full three minutes of stimulation, but a significant difference emerged when only the first minute of exposure was considered, with newborns producing more suck peaks during listening to the structured than the variant sound. This suggests that the effect was most pronounced, and possibly transient, at the beginning of exposure. This transient pattern is compatible with an attentional account of NNS variations in newborns. Valenza et al. (1994) showed that non-nutritive sucking rate transiently decreases at the time the infants’ gaze is captured by a briefly presented peripheral visual cue, interpreting this reduction as a physiological index of the phasic attentional orienting response. Using auditory stimuli, Silvestri et al. (2025, 2026) reported a similar pattern of NNS inhibition in response to auditory stimuli lacking positive socio-emotional prosodic cues (Silvestri et al., 2025) and to sad compared with happy utterances (Silvestri et al., 2026). Applied to the present findings, the variant sound, being less predictable and less temporally regular, and so more complex and unfamiliar in light of the accumulated in-utero rhythmic experience, may have similarly captured attentional resources, resulting in a brief reduction of suck peak rate. In contrast, the structured sound, being more predictable, may not have elicited the same degree of attentional engagement, and may instead have promoted sucking behaviour by allowing attentional resources to remain available for ongoing motor output.

The structured sound also elicited higher suck pressure than the silent baseline, indicating that a temporally regular auditory stream enhances not only the frequency and organization of suck peaks, but also their vigour, again pointing to a facilitatory, organizing effect of auditory regularity on the NNS motor behaviour.

The absence of differences between either auditory condition and the silent baseline for suck burst and peak rates further suggests that the presentation of sounds did not produce a generalized increase or suppression of NNS activity. Instead, the differential response observed between the structured and variant acoustic streams indicates that the temporal characteristics of the auditory input significantly modulated NNS behaviour.

This is further supported by the results of the coherence analyses, which provided more direct evidence that newborns selectively adjusted their sucking behaviour to the temporal structure of the auditory input. NNS-sound coherence was overall higher during exposure to the structured than the variant stream, indicating stronger overall coupling between sucking and the auditory signal when temporal regularities were more pronounced. Importantly, however, this overall increase in coherence was accompanied by a frequency-specific effect: for each auditory stream, coherence was selectively higher at the frequencies distinctive of that sound, indicating fine-grained tracking of stimulus-specific temporal structure rather than a purely stimulus-general modulation of NNS. To the best of our knowledge, this represents the first behavioural evidence of sensorimotor synchronization at birth. This finding challenges the view that precise synchronization to the temporal structure of external stimuli requires substantial postnatal sensorimotor experience (e.g., Provasi et al., 2014; Rocha et al., 2022), and suggests instead that NNS, a motor pattern that is already highly mature at birth, may provide a particularly sensitive window into early rhythm processing and statistical learning abilities.

Notably, frequency-specific coupling was observed not only for the structured stream but also for the variant stream, despite the latter lacking a single dominant periodicity. Importantly, coherence values did not vary across the three minutes of auditory stimulation, indicating that this frequency-specific alignment was already present during the first minute of exposure and did not progressively emerge with prolonged auditory experience. Together, these findings suggest that newborns are able to detect and align their behaviour with temporally relevant regularities even when these are less predictable and do not present a stable periodic pattern. This interpretation is consistent with recent evidence that newborns are sensitive to rhythm-based predictability from birth (Bianco et al., 2026), and echoes recent findings in adults showing that listeners can extract exploitable temporal regularities from non-strictly periodic sequences, effectively “finding rhythm” in randomly timed sounds (van der Werff et al., 2025). Such sensitivity to temporal regularities may therefore represent an early-emerging mechanism for generating temporal predictions under conditions of uncertainty. From an evolutionary perspective, this capacity would be highly adaptive, as natural signals - including speech, vocalizations, and music - are inherently variable, probabilistic, and only partially predictable. Successful interaction with the environment likely depends on mechanisms capable of detecting statistical regularities and generating temporal predictions even when sensory input is characterized by reduced predictability or increased entropy.

How the brain accomplishes this remains an open question. A recent computational account has proposed that synchronization with weakly predictable temporal patterns may emerge through the combined action of Bayesian inference and associative learning (Giomo et al., 2026). Whether such computational mechanisms are already in place at birth, however, remains unknown. Future studies directly manipulating the statistical structure of auditory sequences, will be essential to determine the computational principles that support the earliest forms of sensorimotor synchronization.

In conclusion, this work extends current accounts of early motor adaptation to external rhythms by showing that, already at birth, human infants are capable of frequency-specific auditory-motor synchronization, even in response to highly entropic and weakly predictable auditory sequences. These findings highlight the early capacity of the developing brain to detect and align with temporal regularities in the environment, pointing to the adaptive value of extracting statistical structure and generating predictions from complex and variable sensory inputs.

## Author contributions

**MA**: Conceptualization, Methodology, Investigation, Software, Formal Analysis, Visualization, Data Curation, Validation, Writing - Original Draft Preparation, Writing - Reviews & Editing; **AF:** Software, Formal Analysis, Validation, Visualization, Writing – review & editing; **VS**: Investigation, Writing - Reviews & Editing; **MS**: Software; **LC**: Resources; **MLG**: Resources; **DB**: Software, Formal Analysis, Validation, Writing – review & editing; **VMC**: Conceptualization, Methodology, Validation, Resources, Funding Acquisition, Supervision, Project Administration, Writing – review & editing.

## Acknowledgments

We thank all the families who participated, Paolo Bichisao for constructing the stimuli, and Carlo Toneatto for his assistance with the experiments.

AI technologies (i.e., GPT-5.6 Luna, Claude Opus 5.5) were used during the final stage of manuscript preparation for linguistic editing and polishing of the manuscript’s text, to ensure adherence to English-language grammatical and stylistic standards. The authors critically reviewed and refined the suggested edits and take full responsibility for the integrity and content of the final work. No sensitive, proprietary, or human subject data were provided to or processed by these AI technologies.

This research was supported by a PhD Scholarship from the Ministero dell’Università e della Ricerca to MA, and a grant from the Ministero dell’Università e della Ricerca-NextGenerationEU (PNRR M4.C2.I1.1-Avviso 104/2022, CUP H53D23004090006) to VMC.

## Data availability

The hypotheses, methods and the analysis plan were not preregistered. All stimuli, data and analysis scripts are publicly available at the following link: https://osf.io/3dkxc/overview?view_only=8b740e982f8843d1843499e8af0dbac4

## References

Barlow, S. M., Lee, J., Wang, J., Oder, A., Oh, H., Hall, S., Knox, K., Weatherstone, K., & Thompson, D. (2014). Effects of oral stimulus frequency spectra on the development of non-nutritive suck in pre-term infants with respiratory distress syndrome or chronic lung disease, and preterm infants of diabetic mothers. Journal of Neonatal Nursing, 20(4), 178–188. 10.1016/j.jnn.2013.10.005

Barlow, S. M., Liao, C., Lee, J., et al. (2023). Spectral features of non-nutritive suck dynamics in extremely preterm infants. Pediatric Medicine, 6, 23. 10.21037/pm-21-91

Bianco, R., Tóth, B., Bigand, F., Nguyen, T., Sziller, I., Háden, G. P., Winkler, I., & Novembre, G. (2026). Human newborns form musical predictions based on rhythmic but not melodic structure. PLOS Biology, 24(2), e3003600.

Bobin-Bègue, A., Provasi, J., Marks, A., & Pouthas, V. (2006). Influence of auditory tempo on the endogenous rhythm of non-nutritive sucking. European Review of Applied Psychology, 56(4), 239–245.

Drake, C., Jones, M. R., & Baruch, C. (2000). The development of rhythmic attending in auditory sequences: Attunement, referent period, focal attending. Cognition, 77(3), 251–288. 10.1016/S0010-0277(00)00106-2

Giomo, D., Mancinelli, F., Ravignani, A., & Bueti, D. (2026). Modeling human synchronization to rhythmic patterns with varying statistical regularities. Acta Psychologica, 266, 1–13.

Green, P., & MacLeod, C. J. (2016). SIMR: An R package for power analysis of generalized linear mixed models by simulation. Methods in Ecology and Evolution, 7(4), 493–498. 10.1111/2041-210X.12504

Ladányi, E., Persici, V., Fiveash, A., Tillmann, B., & Gordon, R. L. (2020). Is atypical rhythm a risk factor for developmental speech and language disorders? WIREs Cognitive Science, 11(5), e1528. 10.1002/wcs.1528

Liao, C., Rosner, J. L., Maron, D., Song, D., & Barlow, S. M. (2019). Automatic nonnutritive suck waveform discrimination and feature extraction in preterm infants. Computational and Mathematical Methods in Medicine, 2019, 1–12. 10.1155/2019/7496591

Nguyen, T., Bigand, F., Reisner, S., Koul, A., Bianco, R., Markova, G., … Novembre, G. (2026). Development of auditory and spontaneous movement responses to music over the first postnatal year. eLife, 14, RP107088.

Norcia, A. M., Appelbaum, L. G., Ales, J. M., Cottereau, B. R., & Rossion, B. (2015). The steady-state visual evoked potential in vision research: A review. Journal of Vision, 15(6), 4. 10.1167/15.6.4

Perugini, M., Gallucci, M., & Costantini, G. (2018). A practical primer to power analysis for simple experimental designs. International Review of Social Psychology, 31(1), 20. 10.5334/irsp.181

Poore, M., Zimmerman, E., Barlow, S. M., Wang, J., & Gu, F. (2008). Patterned orocutaneous therapy improves sucking and oral feeding in preterm infants. Acta Paediatrica, 97(7), 920–927.

Popescu, E. A., Popescu, M., Wang, J., Barlow, S. M., & Gustafson, K. M. (2008). Non-nutritive sucking recorded in utero via fetal magnetography. Physiological Measurement, 29(1), 127–139.

Provasi, J., Anderson, D. I., & Barbu-Roth, M. (2014). Rhythm perception, production, and synchronization during the perinatal period. Frontiers in Psychology, 5, 1048. 10.3389/fpsyg.2014.01048

Reinold, E. (1979). Identification and differentiation of fetal movements. Contributions to Gynecology and Obstetrics, 6, 29–32.

Rocha, S., & Addyman, C. (2022). Assessing sensorimotor synchronisation in toddlers using the Lookit online experiment platform and automated movement extraction. Frontiers in Psychology, 13, 897230.

Rocha, S., Attaheri, A., Choisdealbha, A. N., Brusini, P., Flanagan, S. A., Mead, N., … Goswami, U. (2021). Infant sensorimotor synchronisation to speech and non-speech rhythms: A longitudinal study. PsyArXiv. https://doi.org/10.31234/osf.io/[verify]

Rossion, B., Torfs, K., Jacques, C., & Liu-Shuang, J. (2015). Fast periodic presentation of natural images reveals a robust face-selective electrophysiological response in the human brain. Journal of Vision, 15(1), 18. 10.1167/15.1.18

Silvestri, V., Polver, S., M. L. Giannì, et al. (2025). Newborns’ attention to socio-emotional prosodic cues from voices. iScience, 28(12), 113939. 10.1016/j.isci.2025.113939

Silvestri, V., Polver, S., Morniroli, D., Giannì, M. L., Gervain, J., & Bulf, H. (2026). First sounds of emotion: Newborns’ sensitivity to emotional prosody and its relation with maternal depressive traits. Infancy, 31(2), e70087.

Standley, J. M. (2000). The effect of contingent music to increase non-nutritive sucking of premature infants. Pediatric Nursing, 26(5), 493.

Thompson, E. C., White-Schwoch, T., Tierney, A., & Kraus, N. (2015). Beat synchronization across the lifespan: Intersection of development and musical experience. PLOS ONE, 10(6), e0128839. 10.1371/journal.pone.0128839

Tillmann, B., Goswami, U., & Moghimi, S. (2026). Rhythm processing across development: Origins, links to language processing, and perspectives for intervention. Annals of the New York Academy of Sciences, 1556(1), e70161.

Valenza, E., Simion, F., & Umiltà, C. (1994). Inhibition of return in newborn infants. Infant Behavior and Development, 17(3), 293–302.

van der Werff, J., Tufarelli, T., Verga, L., & Ravignani, A. (2025). Humans can find rhythm in randomly timed sounds. Royal Society Open Science, 12(8), 1–11.

Winkler, I., Háden, G. P., Ladinig, O., Sziller, I., & Honing, H. (2009). Newborn infants detect the beat in music. Proceedings of the National Academy of Sciences, 106(7), 2468– 2471. 10.1073/pnas.0809035106

Wolff, P. H. (1968). The serial organization of sucking in the young infant. Pediatrics, 42(6), 943–956.

Woodruff Carr, K., White-Schwoch, T., Tierney, A. T., Strait, D. L., & Kraus, N. (2014). Beat synchronization predicts neural speech encoding and reading readiness in preschoolers. Proceedings of the National Academy of Sciences, 111(40), 14559–14564.

Zentner, M., & Eerola, T. (2010). Rhythmic engagement with music in infancy. Proceedings of the National Academy of Sciences, 107(13), 5768–5773. 10.1073/pnas.1000121107

Zimmerman, E., & Foran, M. (2017). Patterned auditory stimulation and suck dynamics in full-term infants. Acta Paediatrica, 106(5), 727–732.

